# Microglia limit lesion expansion and promote functional recovery after spinal cord injury in mice

**DOI:** 10.1101/410258

**Authors:** Faith H. Brennan, Jodie C.E. Hall, Zhen Guan, Phillip G. Popovich

## Abstract

Traumatic spinal cord injury (SCI) elicits a robust intraspinal inflammatory reaction that is dominated by at least two major subpopulations of macrophages, i.e., those derived from resident microglia and another from monocytes that infiltrate the injury site from the circulation. Previously, we implicated monocyte-derived macrophages (MDMs) as effectors of acute post-injury pathology after SCI; however, it is still unclear whether microglia also contribute to lesion pathology. Assigning distinct functional roles to microglia and MDMs in vivo has been difficult because these CNS macrophage subsets are morphologically and phenotypically similar. Here, to characterize the role that microglia play in experimental models of thoracic spinal contusion or lumbar crush injury, mice were fed vehicle chow or chow laced with a CSF1R receptor antagonist, PLX5622. Feeding PLX5622 depletes microglia. In both groups, spontaneous recovery of hindlimb motor function was evaluated for up to 8 weeks post-SCI using open-field and horizontal ladder tests. Histopathological assessment of intraspinal pathology was assessed in 8 week post-injury tissues. In both SCI models, microglia depletion exacerbated lesion pathology and impaired spontaneous recovery of hind limb function. Notably, the loss of microglia prevented astroglial encapsulation of the lesion core, which was associated with larger lesions, enhanced demyelination and neuron loss and a larger inflammatory response that was dominated by monocyte-derived macrophages. The neuroprotective and healing properties of microglia become obvious in the subacute phases of recovery; microglia depletion up to 7 days post-injury (dpi) had no apparent effect on recovery while delayed depletion from 8-28dpi exacerbated lesion pathology and significantly impaired functional recovery. These data suggest that microglia have essential tissue repair functions after SCI. Selective enhancement of microglial activities may be a novel strategy to preserve tissue and promote recovery of function after neurotrauma.

## Introduction

Abundant macrophage presence is a well-documented neuropathological feature of clinical and experimental spinal cord injury (SCI) (Beck, et al., 2010, Fleming, et al., 2006, Kigerl, et al., 2006, Sroga, et al., 2003). Intraspinal macrophages have diverse roles that can drive events on the spectrum between promoting repair and exacerbating tissue damage (Gensel, et al., 2009, Kigerl, et al., 2009, Shechter, et al., 2009, Shechter, et al., 2013). The function of macrophages depends on several factors including the injury nature and severity, timing and location of activation, and stochastic binding of macrophage receptors by ligands in the injury microenvironment (reveiwed by Brennan and Popovich, 2018). Adding further complexity, intraspinal macrophages are a mixed population derived from blood-borne monocytes and resident CNS microglia.

Early studies showed that intraspinal recruitment of monocyte-derived macrophages (MDMs) peaks at 7-14 d post-injury (dpi), and they persist indefinitely in the gray matter and subpial white matter (Kigerl, et al., 2006, Popovich and Hickey, 2001, Sroga, et al., 2003). Although MDMs can be reprogramed to have reparative functions (Gensel, et al., 2009, Gensel, et al., 2015, Kroner, et al., 2014), a large body of evidence indicates that a disproportionate, predominantly pro-inflammatory MDM response drives poor outcomes from SCI; depleting MDMs or limiting their recruitment improves injury outcomes (Blight, 1994, Blomster, et al., 2013, Brennan, et al., 2015, Fleming, et al., 2008, Geremia, et al., 2012, Giulian and Robertson, 1990, Gris, et al., 2004, Iannotti, et al., 2011, Kigerl, et al., 2009, Lee, et al., 2011, Popovich, et al., 1999). Whether MDM depletion enhances recovery in part by allowing reactive microglia to play a lead role in the recovery process is unknown.

Microglia originate from myeloid lineage precursors in the mesodermal yolk sac and comprise 5-12% of cells in the CNS (Alliot, et al., 1999, Ginhoux, et al., 2010, Kierdorf, et al., 2013). Resting microglia are sentinels of the CNS, using ramified processes to interact with and modulate neighboring blood vessels, neurons, and other glia (Kettenmann, et al., 2011). Upon sensing injury, microglia adopt an amoeboid morphology, proliferate, migrate, and phagocytose cellular debris (Kreutzberg, 1996). Release of neurotrophic factors and removal of debris by activated microglia could promote regeneration of severed axons. Conversely, production of cytokines and free radicals might promote neurotoxicity. Whether microglia act in concert with MDMs or have distinct functions in SCI has been a difficult question to address, since these cells share an identical morphology and express similar phenotypic markers *in vivo* (Guillemin and Brew, 2004).

The identification of microglia-specific molecular signatures, including expression of P2RY12 and TMEM119, has led to the development of antibodies that can distinguish microglia from MDMs *in vivo* (Bennett, et al., 2016, Butovsky, et al., 2014). Moreover, the development of orally bioavailable CSFR1 antagonists enables targeted ablation of microglia in rodent models (Acharya, et al., 2016, Dagher, et al., 2015, Elmore, et al., 2014, Hilla, et al., 2017, Liddelow, et al., 2017). Less than 1% of microglia are detectable in brains of mice on a constant diet containing CSF1R antagonist, while numbers of peripheral macrophages, other immune cells, and overall brain morphology remain intact (Dagher, et al., 2015, Elmore, et al., 2015, Elmore, et al., 2014). Combined, these new experimental tools provide an unprecedented opportunity to define the role that microglia play in regulating outcomes after SCI.

In the present study we took advantage of a specific CSFR1 antagonist, PLX5622, to characterize the role of microglia in mouse models of moderate (75 kdyne contusion) and severe (complete crush) SCI. Our data indicate that microglia are essential for coordinating repair and recovery after SCI. Specifically, without microglia the axon and myelin pathology is exacerbated, pathological MDM infiltrates are enhanced, astrocyte- and NG2-mediated glial scar formation is disrupted and spontaneous recovery of function is impaired. Ongoing studies will define the mechanisms by which microglia coordinate intercellular repair cascades. By doing so, it may be possible to enhance or co-opt microglia functions to accelerate or improve naturally occurring mechanisms of CNS repair to improve clinical SCI outcomes.

## Methods

### Mice and experimental groups

All surgical and postoperative care procedures were performed in accordance with The Ohio State University Institutional Animal Care and Use Committee. Adult (10-14 week old) female C57BL/6J (WT) mice were purchased from Jackson Laboratories (RRID: IMSR_JAX:000664). Mice were age and weight-matched within experiments. Animals were housed under conventional conditions on a 12 hour light-dark cycle with *ad libitum* access to food and water.

A colony stimulating factor 1 receptor (CSF1R)-antagonist, PLX5622 (1200 mg/kg p.o; 290 ppm, Plexxikon)\ was used to pharmacologically deplete microglia. PLX5622 is an orally bioavailable, blood-brain barrier permeable compound that specifically inhibits CSF1R tyrosine kinase activity with 50-fold selectivity over four related kinases (Dagher, et al., 2015). Animals were randomly assigned to cages then cages were randomly assigned to a diet using QuickCalcs (GraphPad software). Animals within a cage received the same diet (i.e. Vehicle or PLX5622). To ensure experimenter blinding during behavioural testing, mice were acclimated to diet-free cages before testing, and these cages were coded (e.g. ‘A’, ‘B’ ‘C’ ‘D’) prior to testing. Experimenters were also blinded to diet group for histology, tissue processing, image acquisition and data analysis by coding the tissue. For dosing studies in naïve (uninjured) mice, mice received PLX5622 diet continuously for 3, 7, 14, or 21 days. For drug washout studies, mice received PLX5622 diet for 14 days, followed by 3 or 7 days of Vehicle diet. Control mice received Vehicle diet for 21 days. For initial contusion SCI studies (35 days post-injury (dpi) endpoint), mice continuously received either Vehicle or PLX5622 diet for 7 weeks; i.e., from -14 dpi to 35 dpi. For contusion SCI studies manipulating timing of microglia depletion (56 dpi endpoint), mice were randomized into groups according to Table 1.

**Table 1.** Feeding schedules and injury biomechanics for different experimental cohorts

| Group | Force (kdyne) | Displacement (µm) | n |
| --- | --- | --- | --- |
| **Experiment 1: 35 dpi endpoint** |  |  |  |
| Vehicle from -14-35 dpi | 78.86 ± 0.83 | 561.6 ± 20.24 | 7 |
| PLX5622 from -14-35 dpi | 76.86 ± 0.59 | 554.1 ± 24.99 | 7 |
| **Experiment 2: 56 dpi endpoint** |  |  |  |
| Vehicle from -14-56 dpi | 78.86 ± 1.28 | 574.1 ± 21.44 | 7 |
| PLX5622 from -14-7 dpi | 78.38 ± 0.96 | 550.8 ± 21.09 | 7 |
| PLX5622 from 8-28 dpi | 77.33 ± 0.95 | 570.2 ± 14.15 | 8 |
| PLX5622 from 29-56 dpi | 76.86 ± 0.63 | 584.3 ± 22.59 | 7 |
| **Experiment 3: 14 dpi endpoint** |  | **Crush duration** |  |
| Vehicle from -14-14 dpi |  | 2 sec | 6 |
| PLX5622 from -14-14 dpi |  | 2 sec | 6 |

### Spinal cord injury and post-operative care

Mice were anesthetized using a cocktail of ketamine (80 mg/kg; i.p.) and xylazine (10 mg/kg; i.p.). For spinal contusion injuries, the ninth thoracic vertebra (T9) was identified based on anatomical landmarks and the lamina removed (Harrison, et al., 2013). Mice received a moderate (75 kdyne) contusion SCI using the Infinite Horizons impactor (Precision Systems and Instrumentation, LLC.) (Scheff, et al., 2003). The mean applied force and tissue displacement for each experimental groups are shown in Table 1. There were no differences in injury parameters between experimental groups.

For crush injuries, the T11 lamina was removed (Harrison, et al., 2013). A complete crush injury was performed at the first lumbar (L1) spinal level by inserting #4 Dumont forceps (Fine Science Tools #11241-30, with tip width narrowing from 0.4-0.2 mm) 2 mm ventrally into the vertebral column on both sides of the spinal cord then laterally compressing the spinal cord for 2s without breaking the dura (Faulkner, et al., 2004, Herrmann, et al., 2008). Injury level and dura integrity were confirmed at post-mortem dissection and using histology.

After SCI, the muscle was closed with 5.0 polyglactin dissolvable sutures, and the skin was closed with wound clips. Animals were injected with saline (2 ml, s.q.) then placed into warmed cages (35°C) until they recovered from anaesthesia. To prevent dehydration and infection, mice were supplemented with daily saline (1-2ml s.q.) and Gentocin (1 mg/kg s.q.) for the first 5 dpi. Because acutely spinal injured mice cannot rear to reach the food hopper in conventional cages, Vehicle or PLX5622 pellets were also placed in the cages for easy access. Bladders were manually voided twice daily for the duration of experiments. Body weight and urinary pH were monitored weekly.

### BrdU administration

To label proliferating cells, mice were injected with the thymidine analog, 5-bromo-2′-deoxyuridine (BrdU) (50 mg/kg i.p. in 0.9% saline; Sigma-Aldrich #10280879001) daily from 1-7 dpi.

### Behavioral testing

#### Open field locomotor testing

Two investigators, blind to treatment, assessed mouse hindlimb function in open-field using the Basso Mouse Scale (BMS) for locomotion (Basso, et al., 2006). BMS testing was performed at -1 (baseline), 1, 3, 7, 10, and 14 dpi, and weekly thereafter until the experimental endpoint (35 or 56 dpi). BMS subscores were also assigned to quantify finer aspects of locomotion.

#### Horizontal ladder task

Three days before SCI, mice were trained to walk along a horizontal ladder as described previously (Cummings, et al., 2007). This task requires mice to navigate across a horizontal ladder with rungs spaced 1.25 cm apart for 40 cm with a mirror below. Mice were video recorded from the side view and the underside view during the task. Mice were motivated to walk along the ladder by placing home cage bedding at the end. A paw falling below the rungs of the ladder during a step in the forward direction was counted as one mistake. The total number of mistakes was averaged across three trials per mouse. Baseline data were recorded beginning 2 d pre-SCI. SCI mice were tested on the horizontal ladder task at 20 dpi, and then weekly thereafter until the experimental end point.

### Tissue Processing

Mice were anesthetized with a lethal mixture of ketamine/xylazine (1.5 × surgical dose) then transcardially perfused with 0.1 M PBS followed by 4% paraformaldehyde. Spinal cords were removed, post-fixed in 4% PFA for 2 h then transferred to 0.1M PBS overnight before being cryoprotected by incubation in 30% sucrose for 48 hours. Spinal cords were blocked into 10 mm segments centered on the impact site with the dorsal columns facing up, then embedded in Tissue-Tek optimal cutting temperature medium (VWR International), and rapidly frozen on dry ice. Tissue sections were cut in series (10 µm thick, 10 slides per series) using a Microm cryostat (HM 505 E) and collected on SuperFrost Plus slides (Thermo Fisher Scientific). Contused thoracic spinal cord segments were cut along the coronal (rostral-caudal) axis. Crushed thoracolumbar spinal cord segments were cut along the horizontal (dorsal-ventral) axis. Slides were stored at -20°C until immunostaining.

### Antibodies and immunostaining

#### Fluorescence immunolabeling

Slides were dried at room temperature (RT) for 2 h, rinsed (0.1M PBS) then blocked with 4% bovine serum albumin and 0.3% Triton X-100 (BP^3+^) in PBS for 1 h at RT. Sections were incubated overnight with primary antibodies (Table 2) at 4°C in humidified chambers. After washing in 0.1 M PBS (3 × 4 mins), sections were incubated with secondary antibodies (Table 2) diluted in 4% bovine serum albumin and 0.1% Triton X-100 (BP^+^) in PBS. Sections were washed again in PBS (3 × 4 mins), then coverslipped with Immumount (Thermo Fisher Scientific). For BrdU labeling, after the initial rinse but before blocking, slides were incubated with 2N HCl for 25 mins at 37°C.

**Table 2.** Immuno-labeling reagents

| *Antigen* | *Host, dilution* | *RRID* | *Vendor, catalog number* |
| --- | --- | --- | --- |
| **Primary antibodies** |  |  |  |
| **BrdU** | Sheep, 1:200 | AB_302944 | Abcam, ab2284 |
| **CD11b** | Rat, 1:200 | AB_321293 | Bio-Rad/AbD Serotec, MCA74G |
| **CD68** | Rat, 1:500 | AB_322219 | Bio-rad/AbD Serotec, MCA1957 |
| **F4/80** | Rat, 1:500 | AB_323279 | Bio-Rad/AbD Serotec, MCA497R |
| **GFAP** | Rabbit, 1:500 | AB_10013382 | Dako, Z0334 |
| **GSTπ** | Rabbit, 1:500 | AB_10778283 | Biorbyt, orb18037 |
| **Iba-1** | Rabbit, 1:500 | AB_839504 | Wako, 019-19741 |
| **MBP** | Rabbit, 1:500 | AB_2313550 | Aves Labs, MBP |
| **NFH** | Chicken, 1:500 | AB_2313552 | Aves Labs, NF-H |
| **NG2** | Rabbit, 1:500 | AB_91789 | Millipore, AB5320 |
| **P2RY12** | Rabbit, 1:1000 | AB_2298886 | Anaspec, AS-55043A |
| **Secondary amplification** |  |  |  |
| **Biotin-chicken IgY** | Goat, 1:1000 | AB_3213506 | Aves Labs, B-1005 |
| **Biotin-rabbit IgG** | Goat, 1:1000 | AB_954902 | Abcam, Ab6720 |
| **Chicken IgY 488** | Goat, 1:500 | AB_2534096 | Thermo Fisher Scientific, A-11039 |
| **Chicken IgY 546** | Goat, 1:500 | AB_2534097 | Thermo Fisher Scientific, A-11040 |
| **Rabbit IgG-488** | Goat, 1:500 | AB_2576217 | Thermo Fisher Scientific, A-11034 |
| **Rabbit IgG-546** | Goat, 1:500 | AB_2534093 | Thermo Fisher Scientific, A-11035 |
| **Rabbit IgG-633** | Goat, 1:500 | AB_2535731 | Thermo Fisher Scientific, A-21070 |
| **Rat IgG 546** | Goat, 1:500 | AB_2534125 | Thermo Fisher Scientific, A-11081 |
| **Streptavidin 488** | 1:250 | AB_2336881 | Thermo Fisher Scientific, S11223 |
| **Flow cytometry antibodies** |  |  |  |
| **CD16/32** | Rat, 1:200 | AB_394657 | BD Biosciences, 553142 |
| **CD11b-PeCy7** | Rat, 1:100 | AB_394491 | BD Biosciences, 552850 |
| **CD11c-BV711** | Hamster, 1:100 | AB_2734778 | BD Biosciences, 563048 |
| **F4/80-BV421** | Rat, 1:50 | AB_2734779 | BD Biosciences, 565411 |
| **Ly6G-PE** | Rat, 1:100 | AB_394208 | BD Biosciences, 551461 |
| **Ly6C-APC** | Rat, 1:100 | AB_1727554 | BD Biosciences, 560595 |

#### Immunoperoxidase labeling

After the initial rinse slides were incubated at RT in methanol containing 6% H_2_O_2_ for 30 mins to quench endogenous peroxidase activity. After washing in 0.1 M PBS (3 × 4 mins), blocking and primary antibody incubations were performed as above. Sections were then incubated with biotinylated secondary antibodies for 1 hour at RT (Table 2). Bound antibody was visualized using Elite-ABC reagent (Vector laboratories) with ImmPACT diaminobenzidine as a substrate (Vector Laboratories Cat #SK-4105). Sections were dehydrated through sequential 2 min incubations in 70%, 70%, 90%, and 100% ethanol solutions, followed by 3 × 2 min incubations in Histoclear. Slides were coverslipped with Permount (Thermo Fisher Scientific).

To visualize myelin, following immunoperoxidase development of neurofilament, slides were rinsed in dH_2_O and immersed in acetone for 8 mins. Samples were then rinsed in dH_2_O before incubation in eriochrome cyanine (EC) for 30 min at RT. Slides were then washed in dH_2_O and differentiated in 5% iron alum and borax ferricyanide for 5-10 mins before dehydration and coverslipping. The injury epicenter was defined visually as the section with the smallest visible rim of spared myelin.

### Analysis of histopathology

#### General histopathology

EC/neurofilament stained sections were digitized using a ScanScope XT scanner (Aperio; OSU Comparative Pathology and Mouse Phenotyping Shared Resource; College of Veterinary Medicine) and ImageScope software (Leica Biosystems). Lesion analysis was performed in ImageJ by inspecting sections 0.2 mm apart across a 4 mm segment extending from 2 mm rostral to 2 mm caudal from the lesion epicenter. Lesioned tissue within this region, delineated by neurofilament staining, was outlined in ImageJ using the polygon selection tool. Lesion volume was calculated by: Σ (area of lesioned tissue per section × distance between sections). Lesion length was the rostral-caudal distance spanned by sections containing lesioned tissue. Three-dimensional reconstructions of lesions were generated with M3D from MCID Elite Software. For lesion epicenter analysis, spared myelin and spared neurofilament were outlined in ImageJ using the polygon selection tool, and expressed as a percentage of the section area.

#### Microglia quantification

For spinal cords cut coronally, sections 0.2 mm apart extending from 2 mm rostral to 2 mm caudal to the epicenter were stained for P2RY12 and developed using immunoperoxidase staining. Sections were digitized using a ScanScope XT scanner (Aperio) and ImageScope (Leica Biosystems). The number of P2RY12^+^ cells were counted using Imaris and expressed as a percentage of the section area, which was outlined in ImageJ.

#### MDM quantification

Spinal sections 0.2 mm apart extending from 2 mm rostral to 2 mm caudal to the epicenter were stained for P2RY12 and the pan macrophage marker F4/80. Sections were imaged using a Leica TCS SP8 confocal microscope. Because MDMs are difficult to individually differentiate within the central lesion core at 35 dpi, the abundance of intraspinal P2RY12^-^F480^+^ MDMs in whole spinal cord sections was expressed as a percentage of the section area. For epicenter region of interest (ROI) analysis, 0.1 mm^2^ boxes were placed in the center of the section and over the ventromedial white matter. The area of P2RY12^-^F480^+^ staining in each ROI was selected using the threshold tool in ImageJ and expressed as a percentage of the ROI area. To assess the size of phagocytically active MDMs in remaining ventral white matter, sections were immunostained with P2RY12, CD68, and myelin basic protein (MBP). Images were captured using a Leica TCS SP8 confocal microscope. The spared ventral white matter was manually outlined as accurately as possible, and the surface area of individual P2RY12^-^CD68^+^ cells in this region were quantified using Imaris (total of 177-395 cells per animal). The average cell area in this region was calculated for each animal.

#### Cell proliferation

Sections were immunostained for GFAP, NG2 and BrdU then imaged using a Leica TCS SP8 confocal microscope. GFAP^+^BrdU^+^ and NG2^+^Brdu^+^ double-labeled cells were counted 0.6 mm rostral and caudal to the epicenter and at the epicenter using Imaris v8.0 (Bitplane Scientific Software). For this, GFAP/BrdU and NG2/BrdU co-localization channels were built based on absolute staining intensity, and touching objects were split. The channel was converted into a surface and thresholds set based on cell area and encapsulation of a BrdU^+^ cell nucleus by GFAP or NG2. NG2^+^ macrophages were excluded using morphological criteria (McTigue, et al., 2001).

#### GFAP intensity analysis

To quantify the intensity of GFAP labeling, horizontal sections from mice with crush SCI were immunostained for GFAP. Sections were imaged using a Leica TCS SP8 confocal microscope and converted to 8-bit format. With the rostral side oriented left, a rectangular box centered over the crush site was drawn on the image. The box was 1.5 mm wide along the rostral-caudal axis and as high as the pia mater; care was taken not to include background pixels. The gray scale profile along the horizontal axis was plotted in ImageJ and the data exported to Microsoft Excel.

#### Quantifying surviving neurons

To estimate the number of neurons in horizontal sections of crushed spinal cords, sections were immunostained for neuronal nuclei (NeuN). Four, 500 µm wide regions of interest (ROI) were overlaid on the section (ROI #1-4), with ROI 1 starting at the lesion edge which was delimited by dense astrogliosis. ROI #2-4 started at the end of the previous ROI, covering a total distance of 2 mm from the lesion edge. NeuN^+^ cell bodies in each ROI were manually counted.

### Flow cytometry

For analysis of circulating leukocyte populations, at the experimental end point blood was collected via cardiac puncture into Microvette 600 µl EDTA capillary tubes. 50µl of blood was mixed with 1ml of red blood cell lysis buffer (0.85% NH_4_Cl in dH_2_O, pH 7.2) then incubated for 5 mins at room temperature. Cells were retrieved via low speed centrifugation (300 × g for 5 mins) and resuspended in DPBS. Samples were incubated with Zombie Green viability dye (BioLegend, #423111) for the exclusion of dead cells. Cells were resuspended in cytometry blocking buffer (0.5% BSA, 2 mM EDTA, in DPBS, pH 7.2), then incubated with rat anti-CD16/32 (1:200; BD Bioscience) for 10 mins on ice to block F_c_ receptors. Cells were incubated with fluorophore-conjugated antibodies (Table 2) for 10 mins on ice. Next, cells were centrifuged (300 × g, 10 mins), washed with cytometry blocking buffer, and resuspended in DPBS with 2 mM EDTA. Unstained, isotype, and fluorescence-minus-one controls were used to establish background staining and autofluorescence. Single stained samples were used as compensation controls. Samples were analyzed using an LSRII flow cytometer and FlowJo software. Compensation was applied to remove spectral overlap. FSC-A vs. SSC-A was used to exclude cell debris and doublets were excluded using FSC-A vs. FSC-H plots.

### Data analysis

GraphpPad Prism was used for data visualization and statistical analyses. BMS and horizontal ladder data were analyzed using two-way repeated-measures ANOVA with Bonferroni post-hoc tests. Two-sided Student’s t-tests were used to compare differences in one variable between two groups, and one-way ANOVA with Bonferroni post-hoc tests were used to compare one variable between three or more groups. A linear regression was used to determine the relationship between tissue pathology and functional recovery. All data were expressed as mean and SEM, with statistical significance determined at p<0.05. Sample sizes were determined *a priori* from historical data using behavioural measures as the primary outcome, with power (1-β) set to 0.8 and α = 0.05.

## Results and Discussion

### CSF1R antagonism depletes microglia in the intact and injured mouse spinal cord

PLX5622 diet depletes brain microglia within 7 d (Dagher, et al., 2015). However, because the brain and spinal cord have different metabolic requirements, vascularization profiles, permeability dynamics, and microglia density (Bartanusz, et al., 2011, Lawson, et al., 1990), we titrated the duration of PLX5622 diet to determine what was optimal for depleting microglia in the intact thoracic spinal cord. We used P2RY12 immunoperoxidase staining to identify microglia (Butovsky, et al., 2014). We confirmed specificity of this antibody in spinal tissue using β-actin^eGFP^>WT bone marrow chimeric mice. Within 3 d of feeding PLX5622 diet to naive mice, spinal microglia were reduced 81% compared to mice fed Vehicle diet (p<0.0001, Fig. 1A, B, H). Microglia numbers were further reduced after 7 d (93.7% reduction), 14 d (99.1% reduction), and 21 d (99.3% reduction) of continuous PLX5622 diet (Fig 1A, C-E, H). When mice were fed PLX5622 diet for 14 d then switched to Vehicle diet, microglia reappeared in the spinal cord within 3 d (80.6% reduced relative to Vehicle diet, p <0.0001), and recovered to baseline levels within 7 d of PLX withdrawal (Fig 1A, F-H). This pattern of depletion and repopulation was reproduced when microglia were quantified using Iba1 or CD11b staining (not shown), suggesting microglia do not simply down-regulate P2RY12 expression as a result of PLX5622 diet. PLX5622 diet for 21 d did not alter the number of NeuN^+^ profiles (p = 0.22; n=3/group), the cross-sectional area of NeuN^+^ cells (p = 0.88; n=3/group), the proportional area (p = 0.39; n=3/group) or intensity (p = 0.67; n=3/group) of GFAP labeling, or the number of GSTπ^+^ cells (p = 0.73; n=3/group). Since myeloid lineage cells other than microglia also express CSF1R (Sasmono, et al., 2003), we evaluated the effect of PLX5622 diet for 21 d on circulating leukocytes in naïve mice. PLX5622 diet did not change blood monocyte (p = 0.21; n=4/group) or granulocyte (p = 0.36; n=4/group) numbers, spleen weight (p = 0.82; n=4/group), or splenic monocyte (p = 0.14; n=4/group) or granulocyte count (p = 0.73; n=4/group).

**Figure 1:**
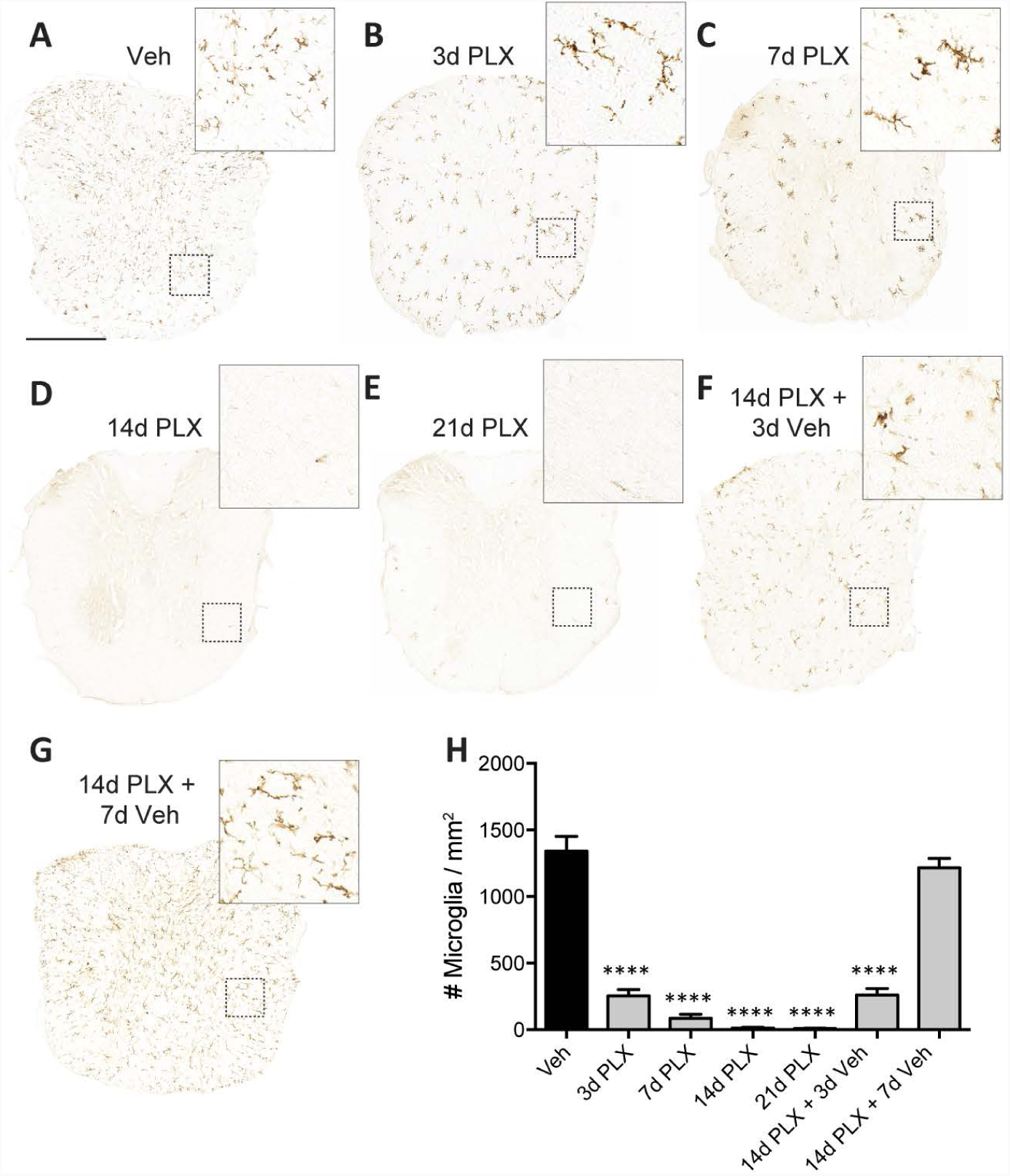
Dynamics of spinal cord microglia depletion and repopulation in naïve mice fed PLX5622 diet (PLX). **A**-**E**: Representative thoracic spinal cord sections stained for P2RY12 showing that compared to vehicle diet (**A**), mice fed PLX5622 for 3 d (**B**), 7 d (**C**), 14 d (**D**), or 21 d (**E**) display a marked reduction in microglia numbers (**H**). Microglia numbers started to repopulate the spinal cord with 3 d of replacing PLX diet with Vehicle diet (**F, H**) and return to control levels within 7 d of diet replacement (**G**, **H**). Scale bar (in A) = 500 µm. One-Way ANOVA with Bonferroni post-hoc tests comparing each group to the Vehicle group; n=3 mice per group; ****p<0.0001.

Next, we assessed the effect of PLX5622 on microglia in the injured mouse spinal cord. Since most microglia in the intact spinal cord were depleted within 14 d of PLX5622 diet, we began PLX5622 diet 14 d prior to contusion SCI with continued feeding of PLX5622 diet until 35 dpi. Control mice were fed a Vehicle diet from -14 dpi to 35 dpi. We assessed the spatial distribution of microglia at 35 dpi using P2RY12 immunostaining. In SCI control mice (Vehicle diet), microglia were sparsely detected in the lesion epicenter; the few present were located in the spared ventrolateral and ventromedial white matter (Fig. 2A). In sections containing injured tissue (∼0.2-0.8 mm rostral and caudal to the epicenter), microglia were absent in the frank lesion area, but prominent amoeboid-shaped microglia clusters were observed at the lesion margins and in the spared white matter (Fig. 2A-D). In sections 1-2 mm from the epicenter, microglia were observed throughout the gray and white matter, with gray matter microglia tending to have a more reactive morphology. In contrast, microglia were significantly reduced (>95%, p<0.0001) throughout the length of the lesion at 35 dpi in mice fed PLX5622 (Fig. 2A-D), as well as in distant cervical and lumbar spinal sections.

**Figure 2:**
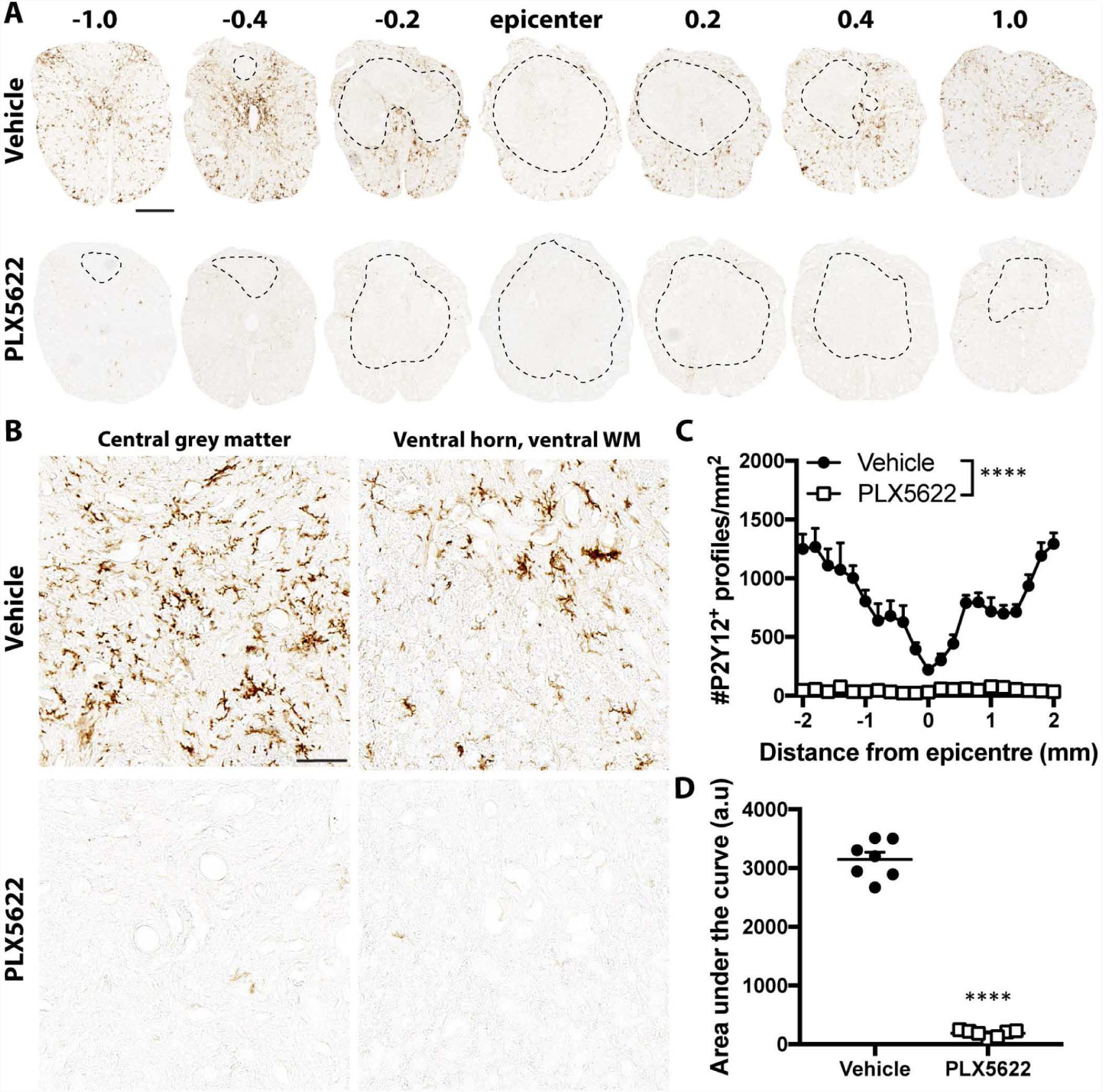
PLX5622 depletes microglia in SCI. **A**: At 35 dpi, intraspinal microglia were identified by immunostaining for P2RY12 (brown). Rostral is left. The frank lesion core is lineated by a black dotted line. In mice fed Vehicle diet (top row), P2RY12^+^ cells were observed at the lesion margins and in the spared white matter, decreasing in abundance toward the epicenter. In mice fed PLX5622 diet (bottom row), P2RY12^+^ cells were rarely observed. Scale bar = 300 µm. **B**: Higher power magnification showing the morphology of P2RY12^+^ cells 2 mm rostral to the lesion epicenter in each group. Scale bar = 50 µm. **C**, **D**: Quantification (**C**) and area under the curve analysis (**D**) confirmed robust depletion of microglia in mice on PLX5622 diet. **C**: Two-Way ANOVA with Bonferroni post-hoc tests; **D**: Student’s two-sided t test; n=7 mice per group; ****p<0.0001.

We also investigated whether PLX5622 affected peripheral cells after SCI. At 35 dpi, mice that had been on PLX5622 diet (for a total of 7 weeks) did not have a reduction in circulating lymphocyte (8.3 ± 0.6 × 10^6^ vs. 9.0 ± 0.5 × 10^6^, p = 0.4038), granulocyte (1.8 ± 0.3 × 10^6^ vs. 1.3 ± 0.2 × 10^6^, p = 0.19) or monocyte numbers (3.0 ± 0.3 × 10^5^ vs. 3.3 ± 0.6 × 10^5^, p = 0.62). Interestingly, at 35 dpi there was a 23% reduction in the proportion of Ly6C^lo^ monocytes in mice fed PLX5622 diet, consistent with previous reports that PLX5622 can alter the proportion of circulating monocyte subsets (Feng, et al., 2017, Valdearcos, et al., 2017). Taken together, these data indicate that PLX5622 depletes microglia in contusion SCI without eliminating peripheral myeloid lineage cells.

### Microglia depletion impairs motor recovery after contusion SCI

The consequences of microglia elimination on functional recovery from T9 contusion SCI were tested by feeding mice Vehicle or PLX5622 diet from -14 d to 35 dpi. Before SCI, all mice displayed normal overground locomotion in the open field, achieving maximum BMS scores of 9. SCI produced near-complete paralysis in all mice at 1 dpi. In mice fed Vehicle diet, locomotor recovery gradually improved and plateaued after 3 weeks (Fig. 3A). By 35 dpi, all mice fed Vehicle diet achieved consistent plantar stepping with 57% (4 of 7) of mice achieving mostly coordinated fore-hindlimb (FL-HL) walking (4 of 7). The remaining mice in this group achieved some FL-HL coordination (43%, 3 of 7). Most mice fed Vehicle diet (85%, 6 of 7) displayed parallel placement of both hind paws at initial contact during the step cycle. For the first week post-injury, mice fed PLX5622 diet recovered similarly to mice fed Vehicle diet; however, after the first week, additional recovery of function was limited in mice fed PLX5622 diet. BMS scores continued to diverge over time, such that mice fed PLX5622 diet were significantly worse than their Vehicle control counterparts at 21, 28, and 35 dpi (Fig. 3A). At 35 dpi, only 29% (2 of 7) of mice fed PLX5622 diet achieved consistent plantar stepping. Only 14% (1 of 7) showed mostly coordinated locomotion, with the remainder achieving only some (43%, 3 of 7) or no (43%, 3 of 7) coordination (Fig. 3A, B). Only 29% (2 of 7) of mice fed PLX5622 diet were able to place both hind paws parallel at initial contact of the step cycle. These behavioral deficits were accompanied by deficits in trunk stability and tail positioning, resulting in significantly worse subscores in mice fed PLX5622 diet from 14-35 dpi (Fig. 3B).

**Figure 3.**
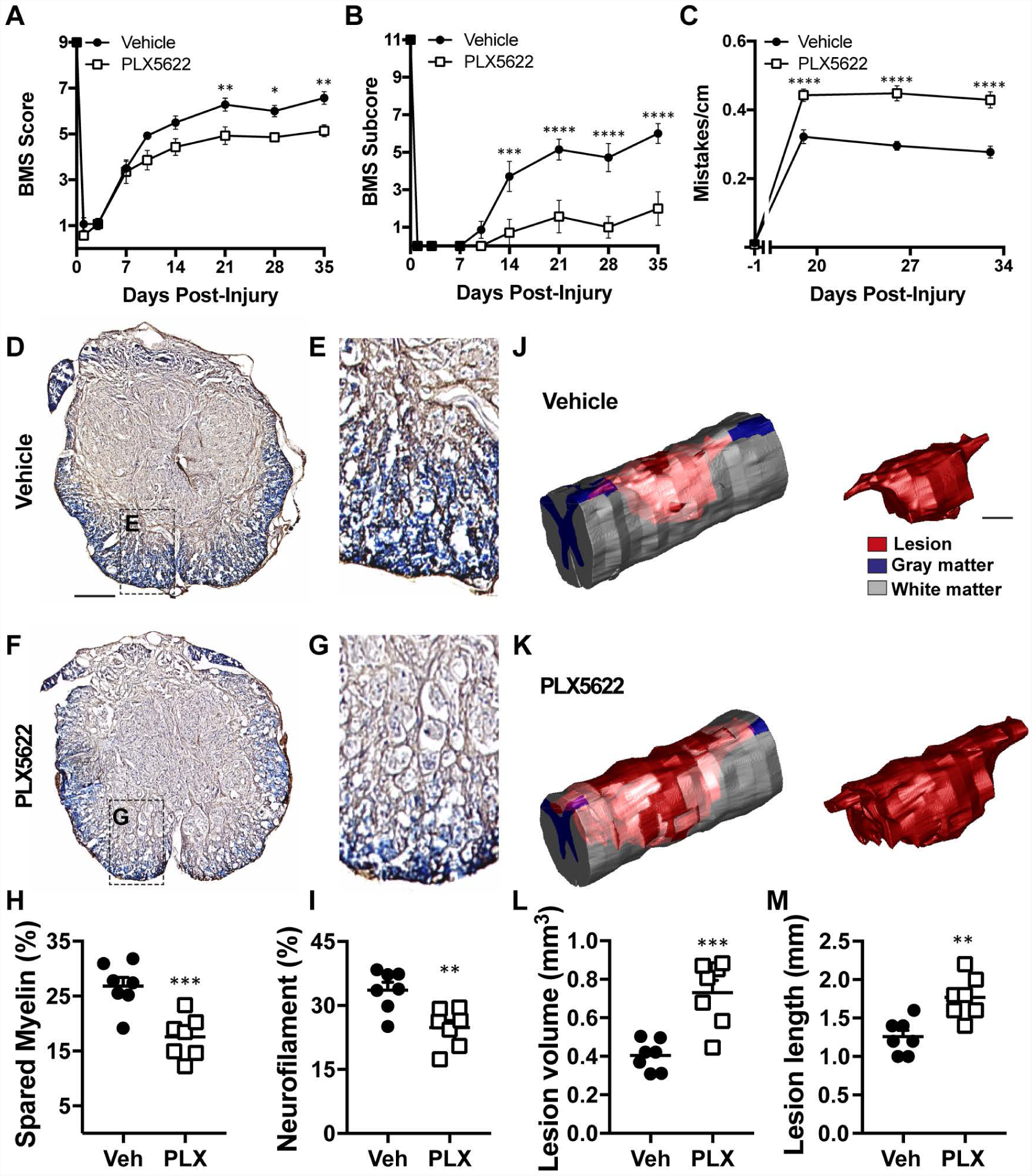
Microglia depletion worsens hindlimb locomotor recovery and increases tissue pathology after contusion SCI. **A**, **B**: BMS scores (**A**) and subscores (**B**) were significantly reduced in mice that received PLX5622 from 21-35 dpi. **C**: The horizontal ladder task revealed that microglia depletion caused mice to made more mistakes from 20 dpi onwards. **A**-**C**: Two-way repeated measures ANOVA with Bonferroni post hoc tests; n=7 per group; *p<0.05; **p<0.01; ***p<0.001; ****p<0.0001. **D**-**G**: EC (blue) and neurofilament (brown) immunostaining indicated diminished presence of white matter and axons as a result of microglia depletion. Scale bar = 200µm. **H**, **I**: Quantitative analysis of the lesion epicenter confirmed reduced myelin sparing (**H**) and axon preservation (**I**) as a result of microglia depletion. **J**, **K**: Three-dimensional reconstructions of the lesion site in mice fed Vehicle diet (**J**) and mice fed PLX5622 diet (**K**). Scale bar = 200µm. **L**, **M**: Both lesion volume (**L**) and lesion length (**M**) was increased by microglia depletion. **H**, **I**, **L**, **M**: Student’s two-sided t tests, n=7 per group; **p<0.01; ***p<0.001.

As a second, independent measure of the effect of microglia depletion on locomotor function, mice were analyzed using the horizontal ladder test. Prior to SCI, mice in both groups had no difficulty completing this task, making few mistakes (Fig. 3C). Although performance on the ladder was impaired after SCI in mice from both groups, mice fed PLX5622 made significantly more stepping mistakes at 20, 27, and 34 dpi (p< 0.0001 vs. Vehicle SCI mice). Taken together, these data show that microglia are essential for achieving optimal spontaneous recovery of locomotor function after SCI, particularly advanced recovery milestones such as stepping, coordination, paw position, and balance.

### Microglia depletion exacerbates tissue loss and causes unusual patterns of neuroinflammation and gliosis after contusion SCI

Microglia depletion caused distinct pathological changes at 35 dpi. Anatomical studies revealed a 35% increase in demyelination (p = 0.001) and a 26% increase in neurodegeneration (p = 0.004) at the lesion epicenter in microglia depleted spinal cords (Fig. 3D-I). Notably, exacerbated myelin and axon pathology were observed in ventral white matter, regions important for overground locomotion in rodents (Loy, et al., 2002, Loy, et al., 2002). Indeed, increased loss of myelin was predictive of poorer BMS scores (r^2^ = 0.64, p=0.0006). Axonal loss was also correlated with worsened locomotor recovery (r^2^ = 0.51, p = 0.0042). Further anatomical analyses revealed that lesion volume (45% p = 0.0006) and lesion length were increased along the rostral-caudal axis (29%, p = 0.0021) in mice fed PLX5622 diet compared to mice on Vehicle (Fig. 3J-M). These data further indicate that microglia are essential for neuroprotection and/or coordinating endogenous recovery processes after SCI.

Given the relationship between neurodegeneration and neuroinflammation (Donnelly and Popovich, 2008, Popovich, et al., 1999), we next explored the effects of microglia depletion on MDM presence. In the epicenter of mice fed Vehicle diet, infiltrating MDM clusters were mostly restricted to the lesion core, fewer MDMs migrate or infiltrate beyond the lesion/white matter interface (Fig. 4A, B). These data are consistent with those described previously in rat and mouse models of SCI (Kigerl, et al., 2006, Mawhinney, et al., 2012, Popovich, et al., 1997, Sroga, et al., 2003, Zhu, et al., 2015). In mice fed PLX5622 diet, MDMs were not confined to the dense core of the lesion epicenter. Rather, large pockets of round phagocytic macrophages extended beyond the lesion boundaries and into spared white matter (Fig. 4C, D). Along the rostral-caudal spinal axis, there was an overall reduction in MDMs as a result of microglia depletion (Fig. 4E). Interestingly, ROI quantification suggested a shift in the localization of MDMs, with mice fed PLX5622 diet having reduced MDM presence in the central core but increased abundance in the ventral white matter (Fig. 4F, G). Co-labeling for CD68, P2RY12 and MBP indicated that phagocytically active MDMs are present in ventral white matter in both groups of mice (Fig. 4H, I). However, CD68^+^ MDMs in ventral white matter of mice on PLX5622 diet had a ‘foamy’ morphology associated with neurotoxicity (Wang, et al., 2015, Zhu, et al., 2017), and were quantitatively larger in size compared to mice fed Vehicle diet (Fig. 4J).

**Figure 4:**
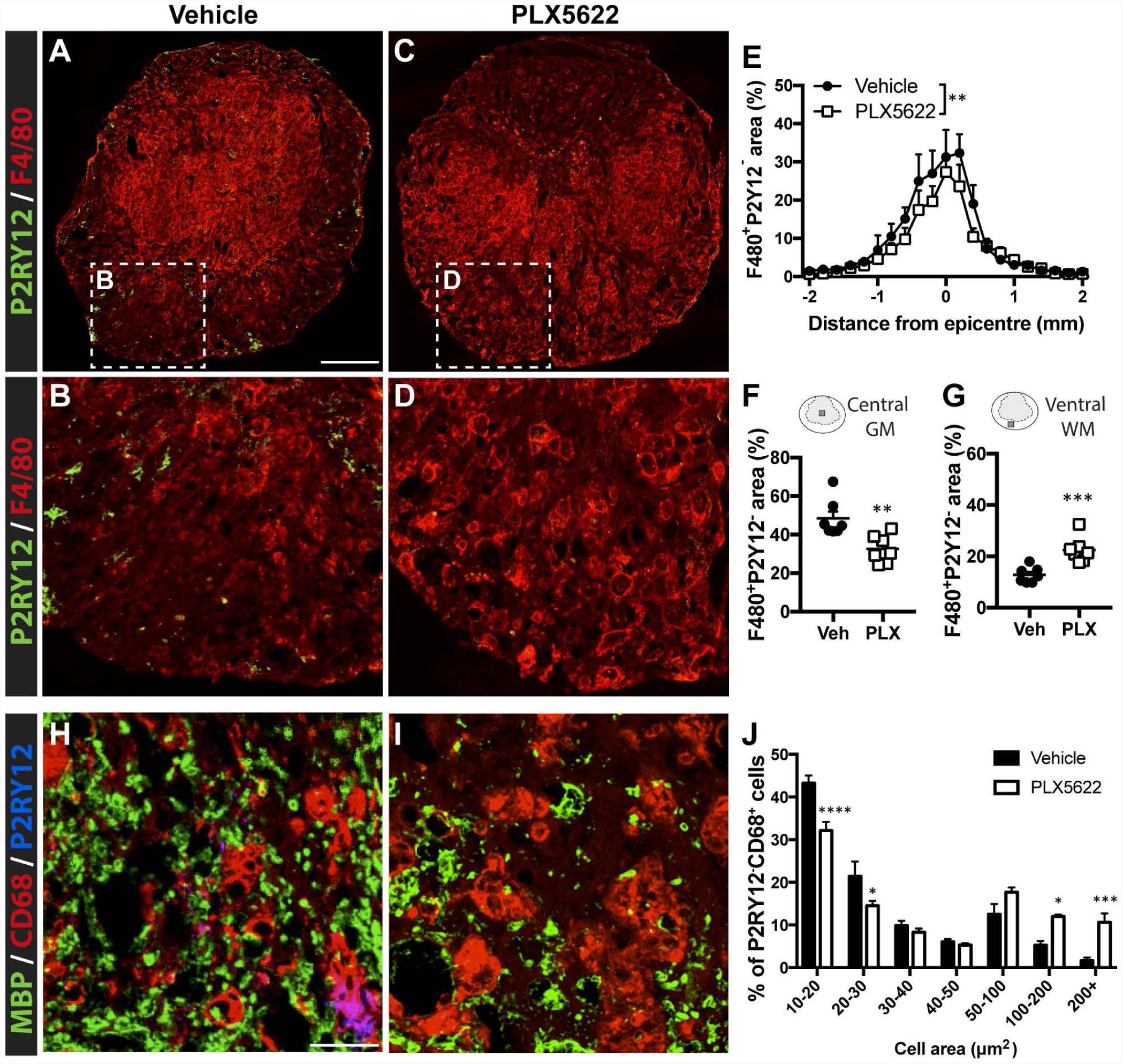
Microglia depletion changes the location and morphology of MDMs in the injured spinal cord at 35 dpi. **A**, **B**: Representative sections of the lesion epicenter from mice given Vehicle diet (**A, B**) or PLX5622 diet (**C, D**). Scale bar = 250 µm. **C**: Quantification of the proportional area of the section occupied by F480^+^P2Y12^-^ cells revealed that microglia depletion causes a reduction in the overall number of MDMs into the injured spinal cord. Two-way ANOVA with Bonferroni post-hoc; n=7 per group, mean + SEM, **p<0.01. **F**, **G**: Region of interest analysis indicated that the proportional area of MDMs was reduced in the central grey matter (GM) (**F**) but increased in the ventral white matter (WM) (**G**) as a result of microglia depletion. **F**, **G**: Student’s two-sided t tests, mean + SEM, n=7 per group; **p<0.01; ***p<0.001. **H**, **I**: Representative images taken from the ventromedial white matter stained for MBP, CD68 and P2RY12 in mice fed Vehicle (**H**) or PLX5622 (**I**) diet. Scale bar = 25 µm. **J**: Area analysis of P2RY12^-^CD68^+^ MDMs in the ventral white matter indicated that microglia depletion was associated with an increase in the size of phagocytically active MDMs. **J**: Two-Way ANOVA with Bonferroni post hoc tests, mean + SEM, n=7 per group; *p<0.05; ***p<0.001; ****p<0.0001.

Because formation of a glial limiting border (i.e., the glial “scar”) is key for the encapsulation of necrotic tissue, containing inflammatory and fibrotic cells from adjacent viable tissue (Faulkner, et al., 2004, Herrmann, et al., 2008, Wanner, et al., 2013), we investigated whether microglia elimination was associated with disrupted glial scar formation. In the lesion epicenter of mice fed Vehicle diet, a robust astrogliotic response was present at the interface between the macrophage-dense lesion core and adjacent spared gray and white matter (White, et al., 2010) (Fig. 5A, B). However, at the lesion epicenter of mice fed PLX5622 diet, large clusters of macrophages were found interspersed between GFAP^+^ cells (Fig. 5C, D). Quantification along the rostral-caudal spinal axis revealed that microglia depletion reduced the proportional area of GFAP^+^ cells at and surrounding the lesion epicenter (Fig. 5E-G).

**Figure 5:**
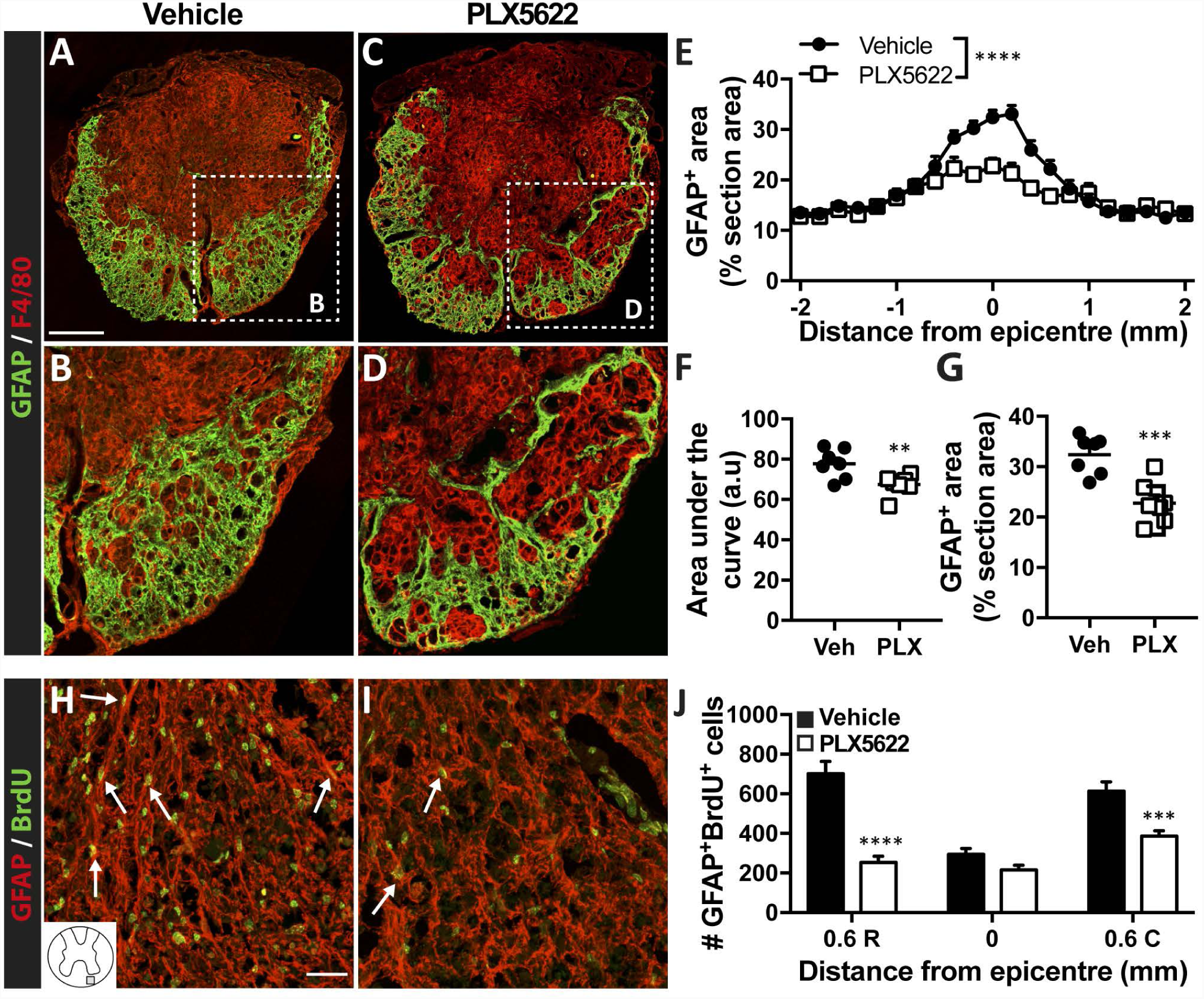
Elimination of microglia reduces glial scar encapsulation of the lesion core. **A**, **B**: Representative images of the lesion epicenter from mice on Vehicle diet (**A**, **B**) and PLX5622 diet (**C**, **D**) mice at 35 dpi immunostained for GFAP and F4/80. High power magnification in **B** and **D** show that mice on PLX5622 diet mice have impaired glial border delineation and many dense clusters of amoeboid-shaped F480^+^ macrophages extending into the spared white matter. Scale bar = 200 µm. **E**-**G**: Quantification of GFAP labeling along the spinal axis (**E**, **F**) and at the lesion epicenter (**G**) confirmed impaired astroglial scar formation as a result of microglia depletion. **E**: Two-Way ANOVA with Bonferroni post-hoc test; **F**, **G**: Two-sided Student’s t tests; n=7 per group; mean + SEM; **p<0.01; ***p<0.001; ****p<0.0001. **H**, **I**: Representative images of GFAP and BrdU staining in the ventral white matter 0.6 mm rostral to the lesion epicenter in mice on Vehicle diet (**H**) or PLX5622 diet (**I**). White arrows point to examples of proliferating astrocytes. **J**: Quantification revealed fewer numbers of proliferating astrocytes rostral and caudal to the lesion epicenter in mice on PLX5622 diet. Two-Way ANOVA with Bonferroni post-hoc test, n=7 per group; mean + SEM; ***p<0.001; ****p<0.0001.

Since the glial scar and demarcation of the lesion is achieved partly through astrocyte hyperplasia (Erturk, et al., 2012), we quantified the number of proliferating (BrdU^+^) astrocytes present at 35 dpi. The number of BrdU^+^GFAP^+^ cells was reduced at the lesion margins in mice fed PLX5622 diet compared to mice fed Vehicle diet (Fig. 5H-J). NG2 cells are also a critical component of glial scar formation (Hesp, et al., 2018). In mice fed Vehicle diet, many NG2^+^ and NG2^+^Brdu^+^ cells were observed at the rostral and caudal margins of the lesion within spared white matter (Fig. 6A-F). Comparatively fewer NG2 glia, both total and proliferating cells, were found in mice fed PLX5622 diet (Fig. 6A-F). Collectively, these data indicate that microglia are essential for coordinating intercellular communication with astrocytes and NG2 glia, i.e., cells that comprise the glial scar and confine inflammatory pathology to the lesion core.

**Figure 6:**
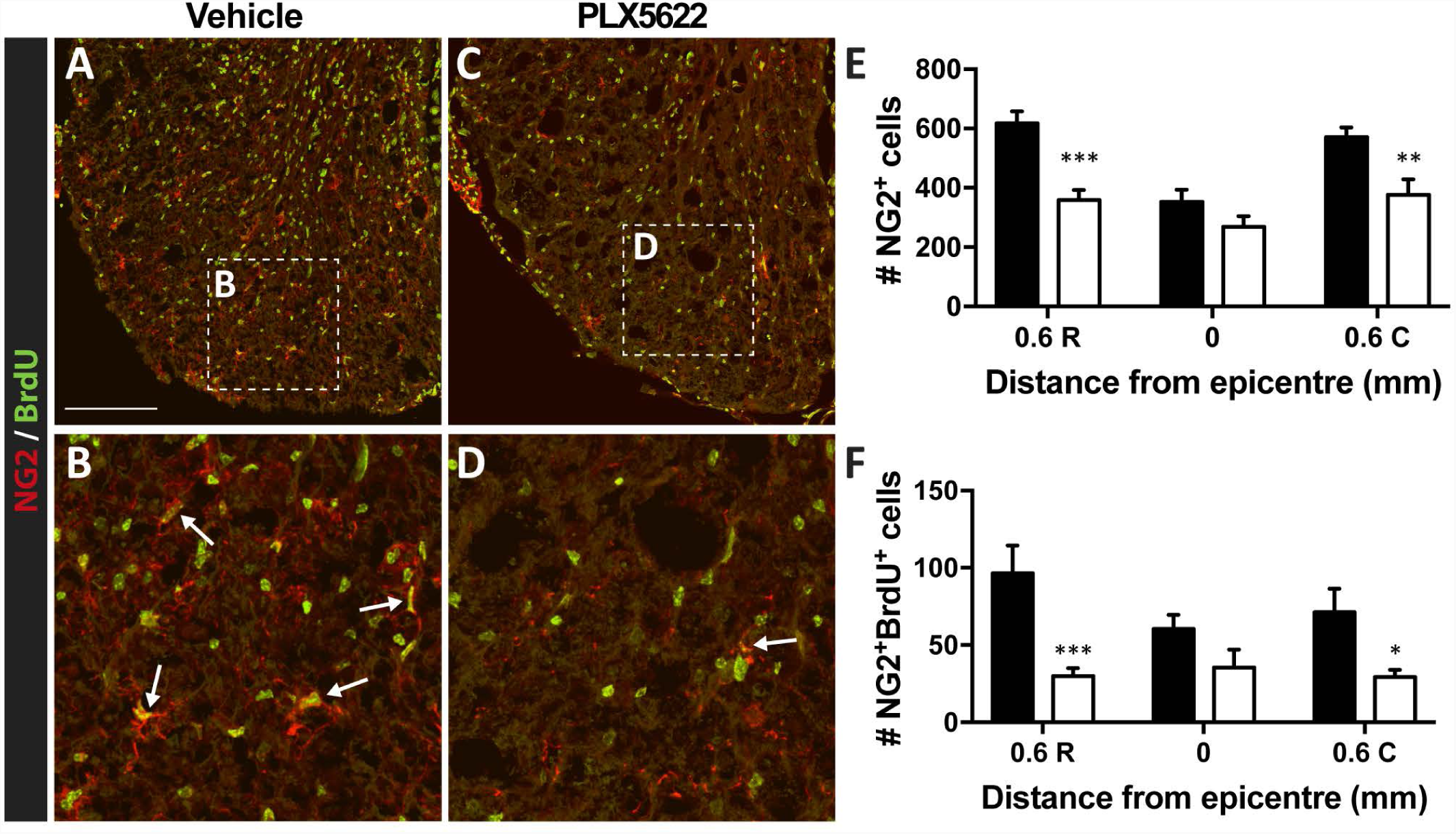
Microglia depletion reduces NG2^+^ glia presence and proliferation. **A**-**D**: Representative images of NG2 and BrdU co-staining in the white matter 0.6 mm rostral to the lesion epicenter from mice on Vehicle diet (**A**, **B**) and mice on PLX5622 diet (**C**, **D**) at 35 dpi. White arrows point to examples of proliferating NG2 glia. Scale bar = 100 µm. **E**-**F**: Quantification of total NG2^+^ cell counts per section (**E**) and NG2^+^BrdU^+^ cell counts per section (**F**) indicated fewer numbers of total and proliferating NG2 glia rostral and caudal to the lesion epicenter in mice with no microglia. **E**, **F**: Two-Way ANOVA with Bonferroni post-hoc tests; n=7 per group; mean + SEM; *p<0.05; **p<0.01; ***p<0.001.

### Microglia are necessary for recovery during subacute and chronic phases of SCI

We next aimed to determine when after SCI, microglia exert their effects on tissue repair and recovery after SCI. To do this we limited microglia depletion to specific post-injury intervals, corresponding with discrete phases of inflammation, lesion expansion and tissue remodeling (Donnelly and Popovich, 2008, Rust and Kaiser, 2017). Mice were fed the PLX5622 diet from -14-7dpi, 8-28dpi, or 29-56 dpi. All mice were fed Vehicle diet when they were not on PLX5622 and were euthanized at 56 dpi. Control mice were fed Vehicle diet for the duration of the study. Consistent with data in Fig. 3A, B, mice continuously fed Vehicle diet showed gradual improvements in locomotor recovery until 21 dpi, after which recovery plateaued (Fig. 7A-C). Analysis of locomotor recovery using BMS or horizontal ladder did not reveal differences between Vehicle control animals and mice fed PLX5622 diet from -14-7dpi (Fig. 7A-C). Conversely, mice fed PLX5622 diet from 8-28 dpi showed deficits in BMS scores from 14 dpi onwards, with functional deficits evident with BMS subscores and the horizontal ladder task from 20-55 dpi (Fig. 7A-C). Mice fed PLX5622 diet from 29-56 dpi showed modest impairments in gross locomotion (detected via BMS and BMS subscores after 29dpi); however, significant deficits were evident when challenged with the horizontal ladder task (Fig. 7A-C).

**Figure 7:**
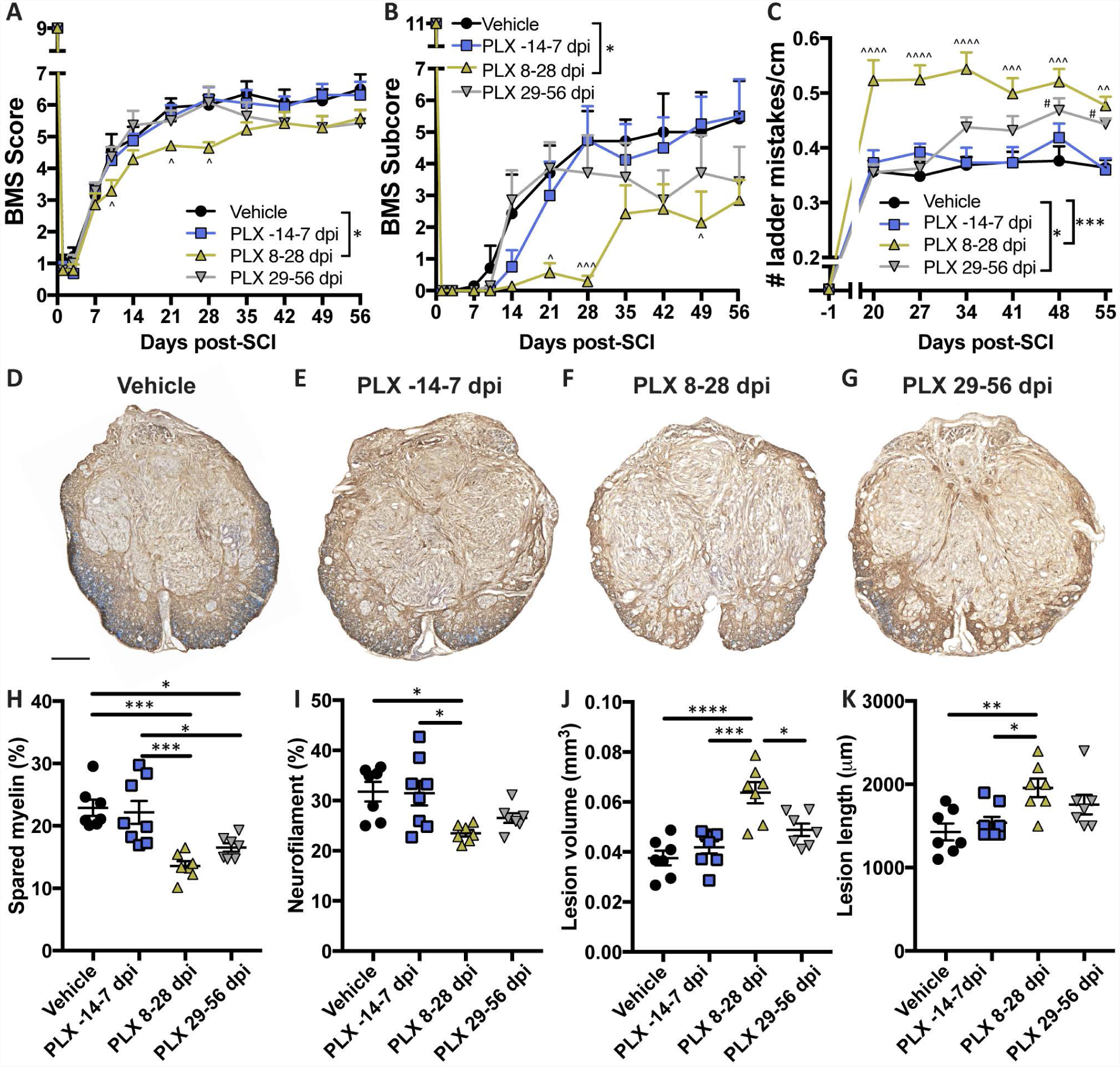
Microglia presence from 8-28 dpi is critical for recovery from SCI in mice. A-C: Microglia depletion during 8-28 dpi worsens BMS scores (**A**), BMS subscores (**B**), and increases the number of mistakes made on the horizontal ladder task (**C**). **A**-**C**: Two-Way repeated measures ANOVA with Bonferroni post hoc tests; n=6-8 per group; mean + SEM; *p<0.05; ***p<0.001; ^p<0.05; ^^p<0.01; ^^^p<0.001; ^^^^p<0.0001 comparing PLX 8-28 dpi to Vehicle; ^#^p<0.05 comparing PLX 29-56 dpi to Vehicle. **D**-**G**: Representative 56 dpi epicenter sections from each group stained for EC/neurofilament. Abnormal ventral white matter pathology is evident in mice on PLX5622 diet from 8-28 dpi, and to a lesser extent in mice on PLX5622 from 29-56 dpi. Scale bar = 200 µm. **H**: Quantification revealed reduced sparing of myelin in mice given PLX5622 from 8-28 dpi or 29-56 dpi. Mice on PLX5622 from 8-28 dpi also showed increased epicenter neurofilament loss (**I**), increased lesion volume (**J**) and lesion length (**K**) compared to mice on Vehicle diet and mice on PLX only from -14-7 dpi. **H**-**K**: One-Way ANOVA with Bonferroni post hoc tests; n=6-8 per group; mean ± SEM; *p<0.05; **p<0.01; ***p<0.001; ****p<0.0001.

Consistent with behavioral data, postmortem histopathological analysis at 56 dpi indicated that there was no difference in myelin/axon sparing, lesion volume or lesion length between mice fed Vehicle diet and mice fed PLX5622 diet from -14-7 dpi (Fig. 7D, E, H-K). However, when microglia were depleted from 8-28 dpi, both demyelination and axon loss were increased, 41% and 26%, respectively, relative to mice fed Vehicle diet (Fig. 7D, F, H-I). Lesion size also increased in mice fed PLX5622 diet from 8-28 dpi (70% increase in volume, 37% increase in length, Fig. 7J, K). Depletion of microglia in the chronic phase of recovery (29-56dpi) also significantly increased demyelination (28% more demyelinated tissue in PLX-fed mice vs. mice fed Vehicle diet, Fig. 7H), and slightly increased axonal loss and lesion size (Fig. 7I-K). Collectively, these data indicate that the microglia effector functions that are most critical for promoting endogenous repair manifest during the subacute period of recovery, i.e. after the first week post-injury.

### Microglia depletion increases inflammatory pathology and impairs glial scar formation in crush SCI

As a second, independent verification that microglia play a pivotal role in coordinating repair processes after SCI and that the effects of the PLX drug were not dependent on either spinal level or injury-severity, we repeated a subset of the above experiments using a bilateral crush injury in the lumber (L1) spinal cord using a previously described injury technique (Faulkner, et al., 2004, Herrmann, et al., 2008). Relative to what happens of spinal contusion injury, glial scar formation is accelerated and complete by 14 dpi after L1 spinal crush injury (Wanner, et al., 2013). In mice fed Vehicle diet, reactive P2RY12^+^ microglia accumulate at the lesion margins at 14 dpi (Fig. 8A, B). Consistent with our titration analyses (Fig. 1) and data in the contused spinal cord (Fig. 2), the number of P2RY12^+^ microglia was reduced along the spinal cord axis in L1 SCI mice fed PLX5622 diet from -14 dpi to 14 dpi (Fig. 8C,D). Quantitative analysis confirmed a significant reduction in the number of microglia at and around the lesion site in lumbar spinal cords of mice fed PLX5622 diet (Fig. 8E).

**Figure 8:**
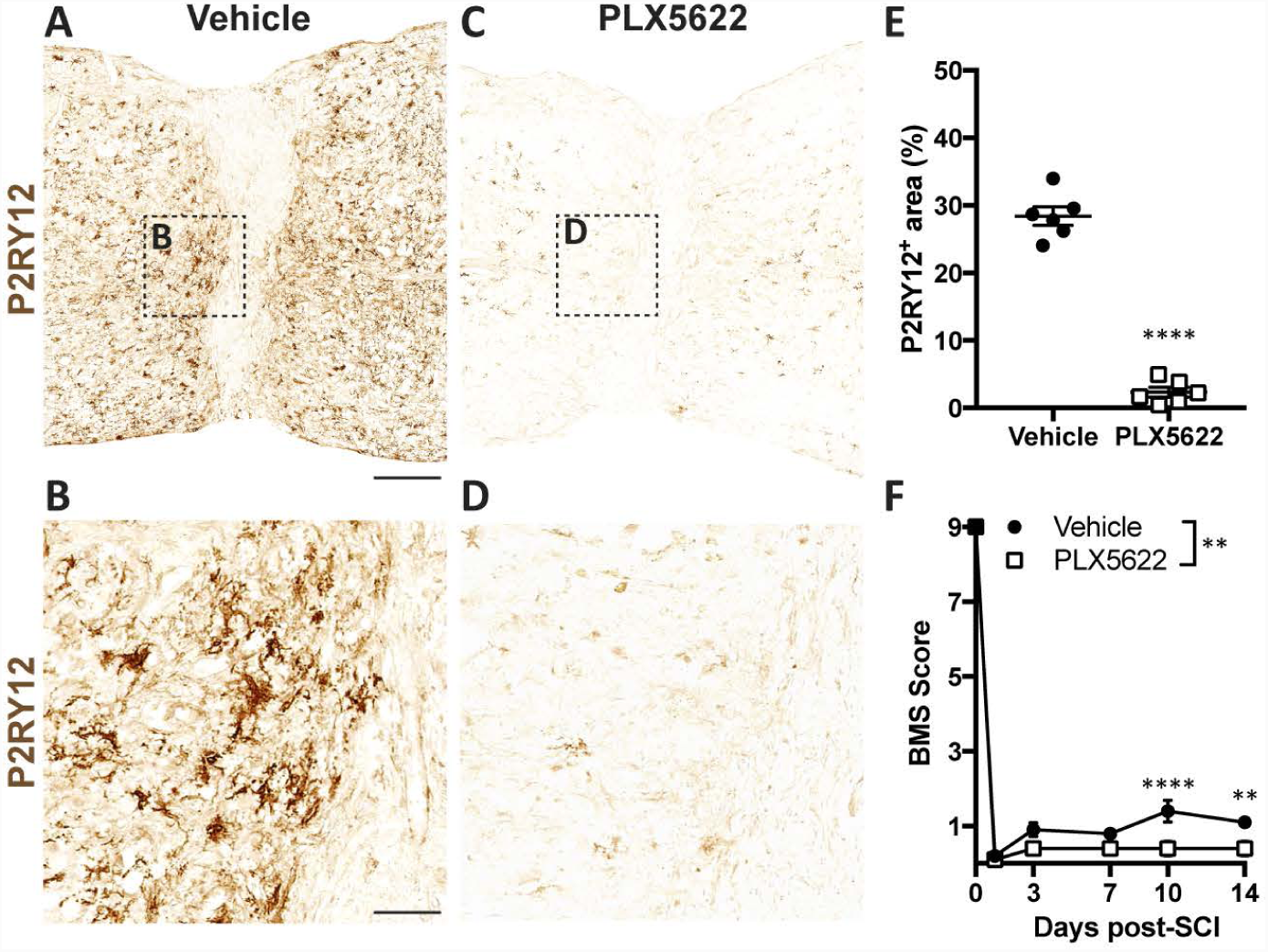
Spinal cord microglia are depleted in mouse L1 crush SCI by PLX5622 diet. **A**-**D**: Representative horizontal sections of spinal cords from injured mice stained with P2RY12. Mice were given Vehicle (**A**, **B**) or PLX5622 diet (**C**, **D**) from 14 d before injury until 14 dpi. Scale bar = 250 µm. **E**: Quantification indicated reduced proportional area of P2RY12 staining in tissue 1.5 mm around in the crush site. Student’s two-sided t-test, n=6 per group; mean ± SEM ****p<0.001. **F**: BMS scoring indicated minimal recovery of hind limb function in both groups of mice, but mice on PLX5622 diet had significantly worse recovery at 10 dpi and 14 dpi compared to mice on Vehicle diet. Two-Way ANOVA with Bonferroni post hoc tests; n=6 per group; mean ± SEM; **p<0.01; ****p<0.0001.

We performed open field BMS testing to assess the functional significance of microglia depletion in lumbar crush SCI. By 14dpi, mice fed Vehicle diet recovered the ability to flex their ankles, often through their entire range of motion (Fig. 8F). This limited recovery is consistent with a severe injury that spares few axons at the injury site (Plemel, et al., 2008). By comparison, mice fed PLX5622 diet did not consistently produce voluntary ankle movements and overall recovery was significantly impaired relative to Vehicle controls at 10 and 14 dpi (Fig. 8F).

As predicted, in spinal cords with microglia (i.e., mice fed Vehicle diet) intense GFAP staining was observed in astrocytes adjacent to the lesion margins (Fig. 9A). This glial-limiting boundary or glial “scar” was morphologically distinct in mice fed PLX5622 diet. Without microglia, astrocytes exhibited a filamentous or lacy phenotype and cells were oriented parallel to the long axis of the spinal cord. This less structured GFAP^+^ interface between lesioned and intact tissue was associated with larger lesions (Fig. 9B,C) and an overall reduction in GFAP immunofluorescence (Fig 9D-G). Without microglia, other parameters of the glial scar also were reduced including the total number of proliferating astrocytes (BrdU^+^GFAP^+^ cells) (Fig. 9H-J) and NG2 glia (BrdU^+^NG2^+^ cells) (Fig. 8K-M). In injured lumbar spinal cord of mice fed PLX5622 diet, demyelination was enhanced (Fig. 10A-C), the overall distribution of F4/80^+^ cells was increased throughout intact spinal parenchyma (Fig. 10D-F), and neuron loss was exacerbated over multiple spinal segments (Fig. 10G-I). Overall, these data replicate those from the thoracic spinal contusion model and illustrate that microglia play a pivotal role in coordinating the formation of the glial scar and limiting the migration of MDMs into intact spinal parenchyma. In a more general sense, microglia are essential for achieving optimal endogenous repair and spontaneous recovery of function after SCI.

**Figure 9:**
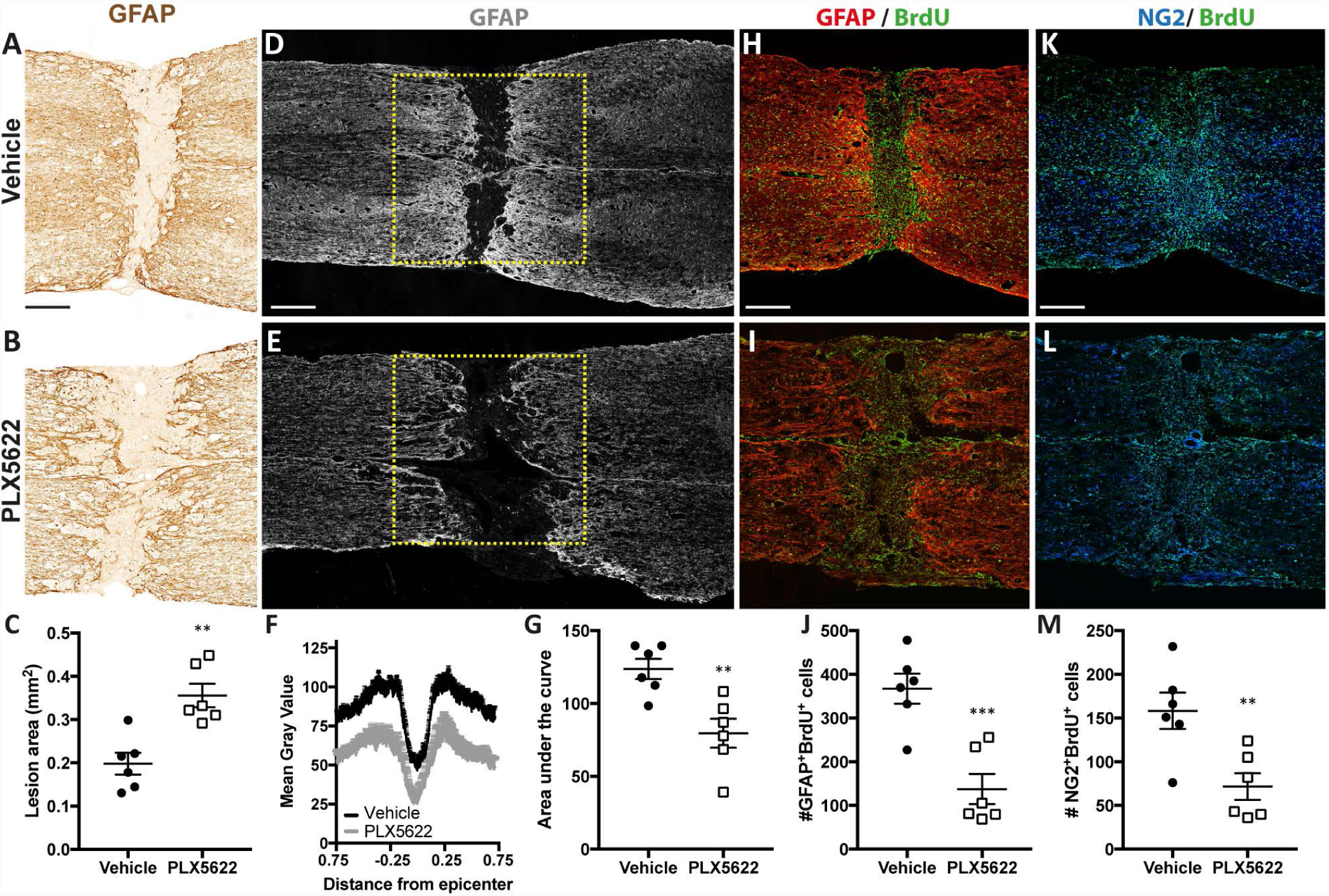
Microglia depletion impairs glial scar formation in L1 crush SCI. Mice were given Vehicle or PLX5622 diet from -14 to 14 dpi and tissue analyzed at 14 dpi. **A**, **B**: Representative horizontal sections of spinal cords immunostained GFAP. Astrocytes formed a dense border in mice on Vehicle diet (**A**), but this response was impaired in mice on PLX5622 diet (**B**). Scale bar = 225 µm**. C**: Quantification indicated larger lesion areas in mice on PLX5622 diet. **D**, **E**: Representative gray scale images of horizontal sections immunofluorescently stained for GFAP. The yellow box indicates the region analyzed in **F** and **G**. Scale bar = 345 µm. **F**, **G**: Mean gray value for the lesion site along the rostral-caudal axis (**F**) and area under the curve analysis (**G**) also indicated reduced GFAP expression in mice on PLX5622. **H**, **I**: Representative horizontal spinal cord sections stained for GFAP and BrdU from mice given Vehicle (**H**) or PLX5622 (**I**) diet. Scale bar = 320 µm **J**: Quantification indicated fewer proliferating astrocytes at the lesion site in the absence of microglia. **K**, **L**: Representative horizontal spinal cord sections stained for NG2 and BrdU from mice given Vehicle (**K**) or PLX5622 (**L**) diet. Scale bar = 320 µm **M**: Quantification indicated fewer proliferating NG2 glia without microglia. **C**, **G**, **J**, **M**: Student’s two-sided t tests; n=6 per group, mean ± SEM; *p<0.05; **p<0.01; ***p<0.001.

**Figure 10:**
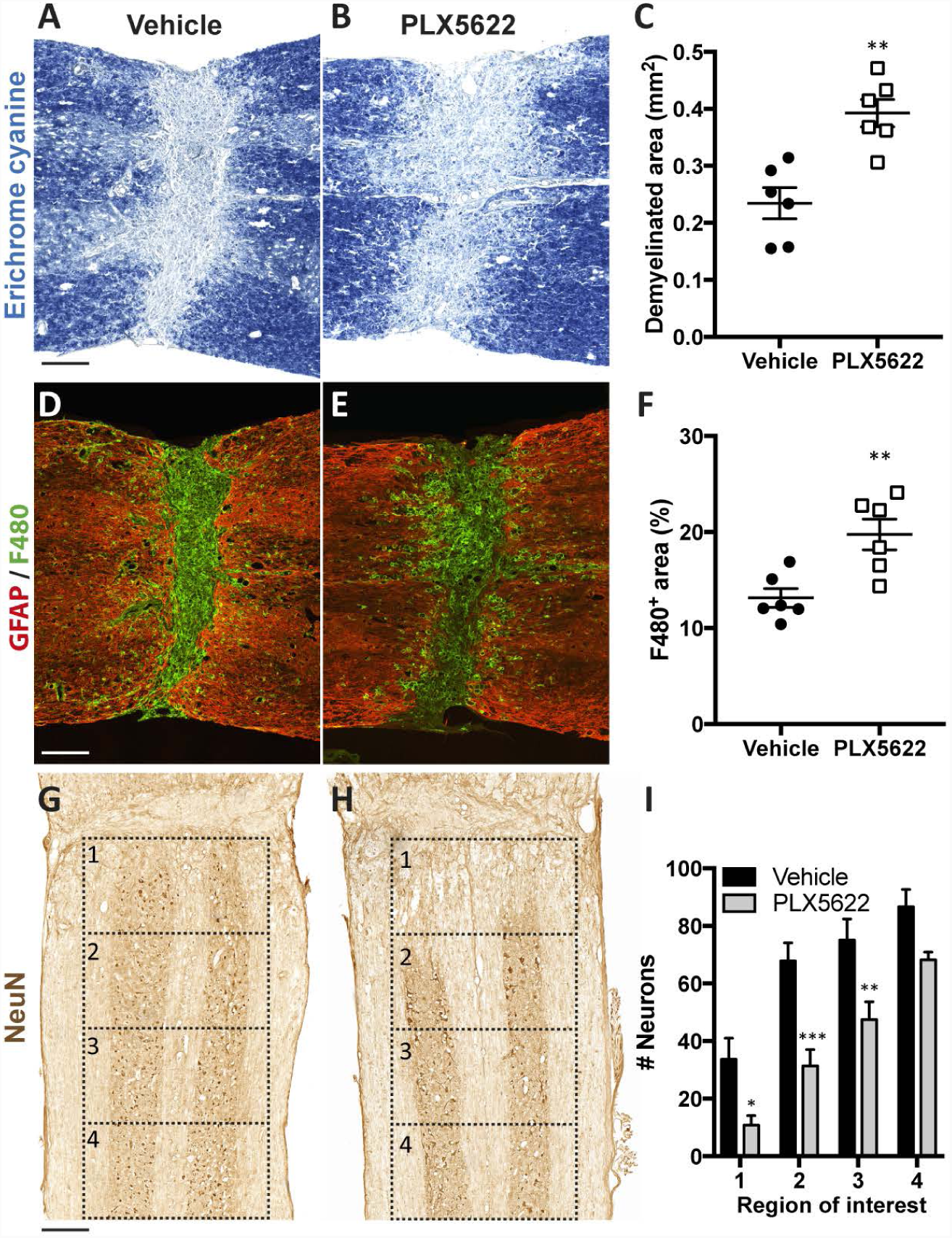
Microglia depletion exacerbates tissue pathology after lumbar crush SCI. Mice were given Vehicle or PLX5622 from -14 to 14 dpi and tissue analyzed at 14 dpi. **A**, **B**: Representative horizontal spinal cord sections stained for EC from mice given Vehicle (**A**) or PLX5622 (**B**). Scale bar = 185 µm. **C**: Quantification indicated enlargement of the demyelinated area in the absence of microglia. **D**, **E**: Representative images of GFAP and F4/80 staining in mice given Vehicle (**D**) or PLX5622 (**E**). Scale bar = 185 µm. **F**: Quantification of the F4/80^+^ area revealed MDMs were not confined to the lesion core in mice on PLX5622. **H**: Representative images showing staining for NeuN at 14 dpi from mice on Vehicle (**G**) or PLX5622 (**H**) diet. Scale bar = 250 µm. **I**: Quantification of NeuN^+^ cell counts indicated reduced numbers of NeuN^+^ cell bodies near the lesion in mice on PLX5622 diet. **C**, **F**: Student’s two-sided t tests; n=6 per group; mean ± SEM **p<0.01. **I**: Two-Way ANOVA with Bonferroni post-hoc tests; n=6 per group; mean + SEM *p<0.05; **p<0.01; ***p<0.001.

Although the role of macrophages in SCI has been explored for the past two decades, the contribution of microglia-derived macrophages in SCI pathology has remained enigmatic. Here, we took advantage of a recently available CSF1R antagonist, PLX5622, to specifically deplete microglia. In doing so, we determined that microglia have an essential protective role after SCI. After both moderate thoracic contusion and severe lumbar crush SCI, elimination of microglia exacerbated motor deficits. This phenotype was accompanied by an abnormal abundance of large, phagocytically active MDMs into otherwise spared tissue, increased demyelination and axonal/neuron loss, and impaired astrocyte- and NG2-dependent encapsulation of the lesion core. Therapies that selectively enhance microglia activity could improve outcomes from SCI and other forms of acquired neurotrauma.

## Acknowledgements

This work is supported by the Craig H. Neilsen Foundation (FHB), The National Institutes of Health (R01NS099532 and R01NS083942 to PGP) and the Ray W. Poppleton Endowment (PGP). The authors thank Andrey Reymar (Plexxikon, Inc.) and Steven Yeung (Research Diets Inc.) for providing Vehicle and PLX5622 diets.

